# Comparative analyses of parasites with a comprehensive database of genome-scale metabolic models

**DOI:** 10.1101/772467

**Authors:** Maureen A. Carey, Gregory L. Medlock, Michał Stolarczyk, William A. Petri, Jennifer L. Guler, Jason A. Papin

**Author notes:** co-corresponding **Contact information:** Maureen A. Carey, Jason A. Papin.

## Abstract

Protozoan parasites cause diverse diseases with large global impacts. Research on the pathogenesis and biology of these organisms is limited by economic and experimental constraints. Accordingly, studies of one parasite are frequently extrapolated to infer knowledge about another parasite, across and within genera. Model *in vitro* or *in vivo* systems are frequently used to enhance experimental manipulability, but these systems generally use species related to, yet distinct from, the clinically relevant causal pathogen. Characterization of functional differences among parasite species is confined to *post hoc* or single target studies, limiting the utility of this extrapolation approach. To address this challenge and to accelerate parasitology research broadly, we present a functional comparative analysis of 192 genomes, representing every high-quality, publicly-available protozoan parasite genome including *Plasmodium, Toxoplasma, Cryptosporidium, Entamoeba, Trypanosoma, Leishmania, Giardia*, and other species. We generated an automated metabolic network reconstruction pipeline optimized for eukaryotic organisms. These metabolic network reconstructions serve as biochemical knowledgebases for each parasite, enabling qualitative and quantitative comparisons of metabolic behavior across parasites. We identified putative differences in gene essentiality and pathway utilization to facilitate the comparison of experimental findings. This knowledgebase represents the largest collection of genome-scale metabolic models for both pathogens and eukaryotes; with this resource, we can predict species-specific functions, contextualize experimental results, and optimize selection of experimental systems for fastidious species.

## Introduction

Malaria, African sleeping sickness, many diarrheal diseases, and leishmaniasis are all caused by eukaryotic single-celled parasites; these infections result in over one million deaths annually and contribute significantly to disability-adjusted life years (Programme 2018; World Health Organization 2018, 2012). In addition, human infectious and related parasites infect domestic and wild animals, resulting in a large reservoir of human pathogens and diseased animal population (Elsheikha and Khan 2011). This combined global health burden makes parasitic diseases a top priority of many economic development and health advocacy groups (Day 2011; May 2007; McCoy et al. 2009). However, effective prevention and treatment strategies are lacking. No widely-used, efficacious vaccine exists for any parasitic disease (*e.g.* (Ghorbani and Farhoudi 2018; Sacks 2014; Dumonteil, Herrera, and Buekens 2019; Pance 2019; Checkley et al. 2015)). Patients have limited treatment options because few drugs exist for many of these diseases, drug resistance is common, and many drugs have stage specificity (*e.g.* (Sparks et al. 2015; Menard and Dondorp 2017; Delves et al. 2012)). Thus, there is a pressing need for novel, effective therapeutics. Beyond the economic constraints associated with antimicrobial development (Simpkin et al. 2017; DiMasi, Hansen, and Grabowski 2003), antiparasitic drug development is technically challenging for two primary reasons: these parasites are eukaryotes and they are challenging to manipulate *in vitro*.

As protozoa, these parasites share many more features with their eukaryotic host than prokaryotic pathogens do. Thus, antiparasitics must target the parasite while minimizing the effect on potentially similar host targets, similar to cancer therapeutics. Enzyme kinetics can be leveraged such that the drug targets the pathogen’s weak points while remaining below the lethal dose for host (Haanstra et al. 2017) or drugs can synergize with the host immune response (*e.g.* Bogdan et al. (1991) and Kumaratilake et al. (1997)). Unique parasite features (*i.e.* signalling cascades as in Zheng (2013) or plastid organelles as in Dahl et al. (2006)) can also be targeted once identified.

Drug target identification and validation are further complicated by experimental challenges associated with these parasites. Many of these organisms have no *in vitro* culture systems, such as *Plasmodium vivax* (malaria) and *Cryptosporidium hominis* (diarrheal disease), or *in vivo* model system, such as *Cryptosporidium meleagridis* (diarrheal disease). Some parasite species have additional unique biology and resultant experimental challenges hindering drug development, such as resistance to genetic modification. For example, *Plasmodium falciparum* (malaria) was considered refractory to genetic modification until recently (Ghorbal et al. 2014; Lee and Fidock 2014). *Entamoeba histolytica* (diarrheal disease) has also been refractory to efficient genetic manipulation, and the genomes of *Leishmania* develop significant aneuploidy under selective pressure (Downing et al. 2011; Sterkers et al. 2012).

Although these challenges may be circumvented with new technology, the use of clinical samples, and reductionist approaches, little data exist relative to that which is available for most bacterial pathogens. Without adequate profiling data (genome-wide essentiality, growth profiling in diverse environmental conditions, etc.), we do not have the knowledge to rationally identify novel drug targets. Untargeted and unbiased screens of chemical compounds for antiparasitic effects have proven useful (if the parasite can be cultured, *e.g.* (Jumani et al. 2018; Chao et al. 2018; Love et al. 2017; Subramanian et al. 2018; Lucantoni et al. 2013)), but this approach provides little information about mechanism of action or mechanisms of resistance development. Typical approaches to study drug resistance, such as evolving resistance to identify mutations in a drug’s putative target, are not possible without a long-term culture system and a relatively well-annotated genome.

As a result of these difficulties (**Figure 1A**), data collected in one organism are frequently extrapolated to infer knowledge about another parasite, across and within genera (**Figure 1B**). *Toxoplasma gondii* is frequently used as a model organism for other apicomplexa due to its genetic and biochemical manipulability (Li and He 2017; Meyer, Caton, and Shapiro 2018; Sidik et al. 2016; Kim and Weiss 2004). Mouse models of malaria (Huang, Pearman, and Kim 2015; Minkah, Schafer, and Kappe 2018) and cryptosporidiosis (Ward 2017; Sateriale et al. 2019) imperfectly represent the disease and/or use different species than the human pathogen. However, the modest characterization of functional differences among parasite species limits the utility of this extrapolation-based approach. Systematic assembly of existing knowledge about parasites and their predicted capabilities could greatly improve the extrapolation-based knowledge transfer by facilitating rigorous *in silico* comparison. Such systems biology approaches (*e.g.* genome-scale metabolic modeling) provides a framework to understand parasite genomes, highlight knowledge gaps, and generate high-confidence data-driven hypotheses about parasite metabolism.

**Figure 1:**
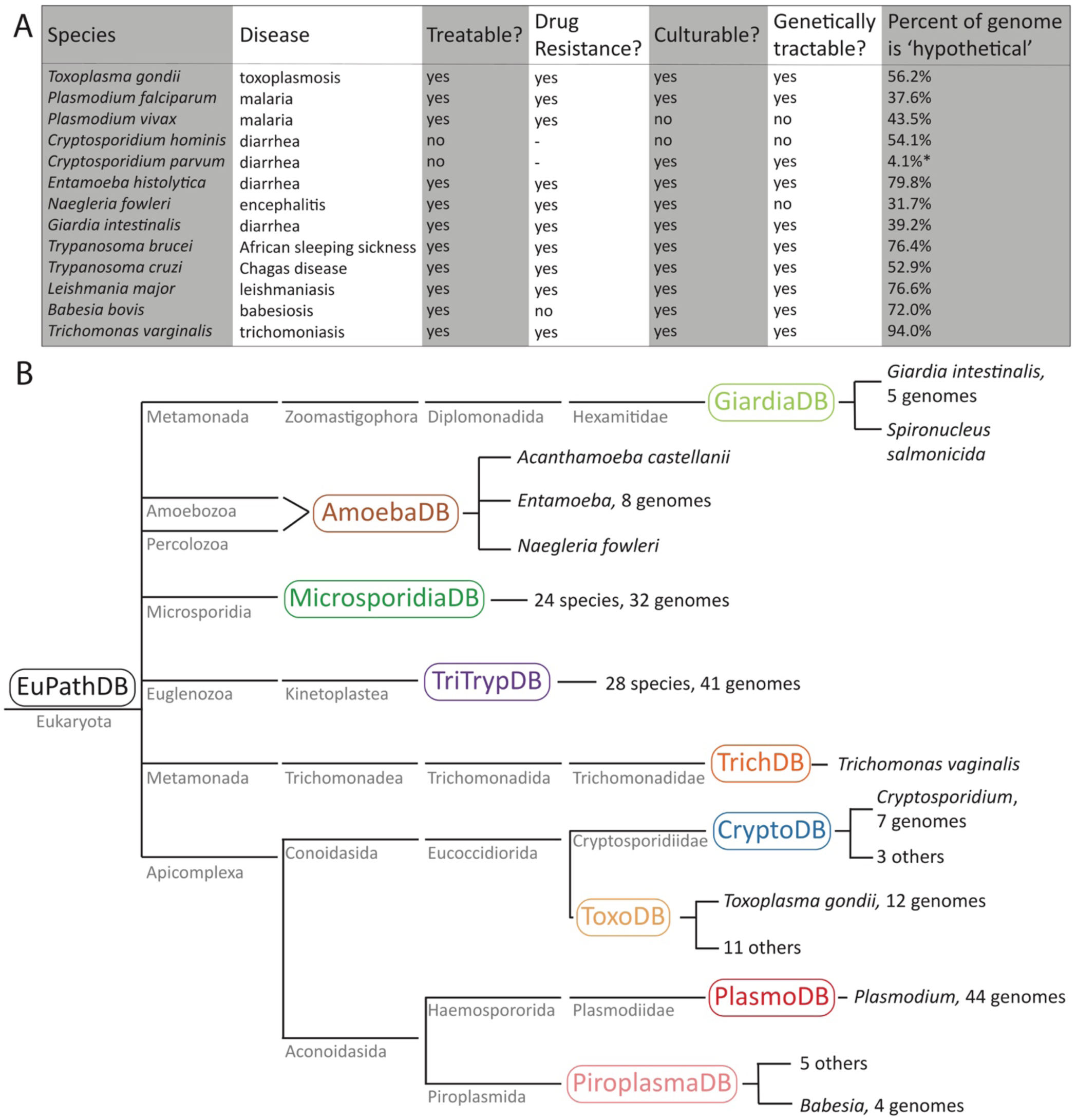
Challenges in parasitology. **(A): Experimental challenges.** Parasites cause important human and animal diseases and have unique biological and experimental challenges that have made interpretation of *in vivo* and *in vitro* data challenging. Several examples are shown. Current treatments and associated observed drug resistance are noted. Many well-studied parasites remain refractory to genetic modification and/or still have poor genome annotation. ‘Uncharacterized’ genes were identified via EuPathDB searches for terms such as ‘uncharacterized’, ‘putative’, ‘hypothetical’, etc. Because each database is heavily influenced by the respective scientific community, some databases such as CryptoDB do not use these terms because the function of so few genes have been validated in the *Cryptosporidium* parasites. Thus, the genomes of the *Cryptosporidium* parasites are mostly hypothetical and proposed functions are only putative; the reported percent of genome that is hypothetical is low for this reason (highlighted by a *). **(B): EuPathDB database.** EuPathDB is the Eukaryotic Pathogens database and serves as a repository for parasite ‘omics data; EuPathDB contains field-specific databases including GiardiaDB, AmoebaDB, MicrosporidiaDB, TriTrypDB, TrichDB, CryptoDB, ToxoDB, PlasmoDB, and PiroplasmaDB (all shown), as well as FungiDB, HostDB, and MicrobiomeDB (*not shown*). Here, a phlogenetic tree of database member parasites is shown. Each EuPathDB sub-database is in a rough phylogenetic grouping, but the parasites on the EuPathDB databases are genetically and phenotypically highly diverse.

Here, we present a parasite knowledgebase, **Para**site **D**atabase **I**ncluding **G**enome-scale metabolic **M**odels (**ParaDIGM**), for this purpose. Metabolic models are built from genomic data and by inferring function to complete or connect metabolic pathways; these models are supplemented with functional genetic and biochemical studies, representing our best understanding of an organism’s biochemistry and cellular biology. The integration of this genomic and experimental evidence into genome-scale metabolic models enables direct comparison of predicted metabolic capabilities in specific contexts, rather than the purely qualitative comparisons that can be performed with traditional genomic approaches. With ParaDIGM, we compare metabolic capacity, gene essentiality, and pathway utilization to better leverage experimentally tractable model systems for the study of eukaryotic parasites and antiparasitic drug development.

## Results

### Building ParaDIGM, a parasite knowledgebase

To build a comprehensive collection of genome-scale network reconstructions representing parasite metabolism, we designed a novel network reconstruction pipeline optimized for eukaryotic organisms (**Figure 2A**). Our pipeline uniquely focuses on the appropriate compartmentalization of biochemical reactions. We applied this pipeline to assemble networks for all publically available reference genomes from parasite isolates representing 119 species (see **Supplemental Information** for link to code and reconstructions). In brief, we obtained 192 high-quality genomes from the parasite genome resource, EuPathDB (Aurrecoechea et al. 2017) to generate a *de novo* reconstruction for each genome (**Figure 2A**, *step 1*). We mapped the protein sequence of all open reading frames against a biochemical database (King et al. 2016) to identify putative metabolic functions via gene-protein-reaction mappings. Reaction compartmentalization was adjusted to maintain each gene-protein-reaction mapping but only with the subcellular compartments relevant for each organism. A large proportion of parasite gene-reaction pairs would otherwise be misassigned or removed from the network due to assignment to an incorrect compartment, due to lack of orthologous and *compartmentalized* reactions in biochemical databases; our pipeline reassigns these erroneous reactions to the cytosol (**Figure 2B**).

**Figure 2:**
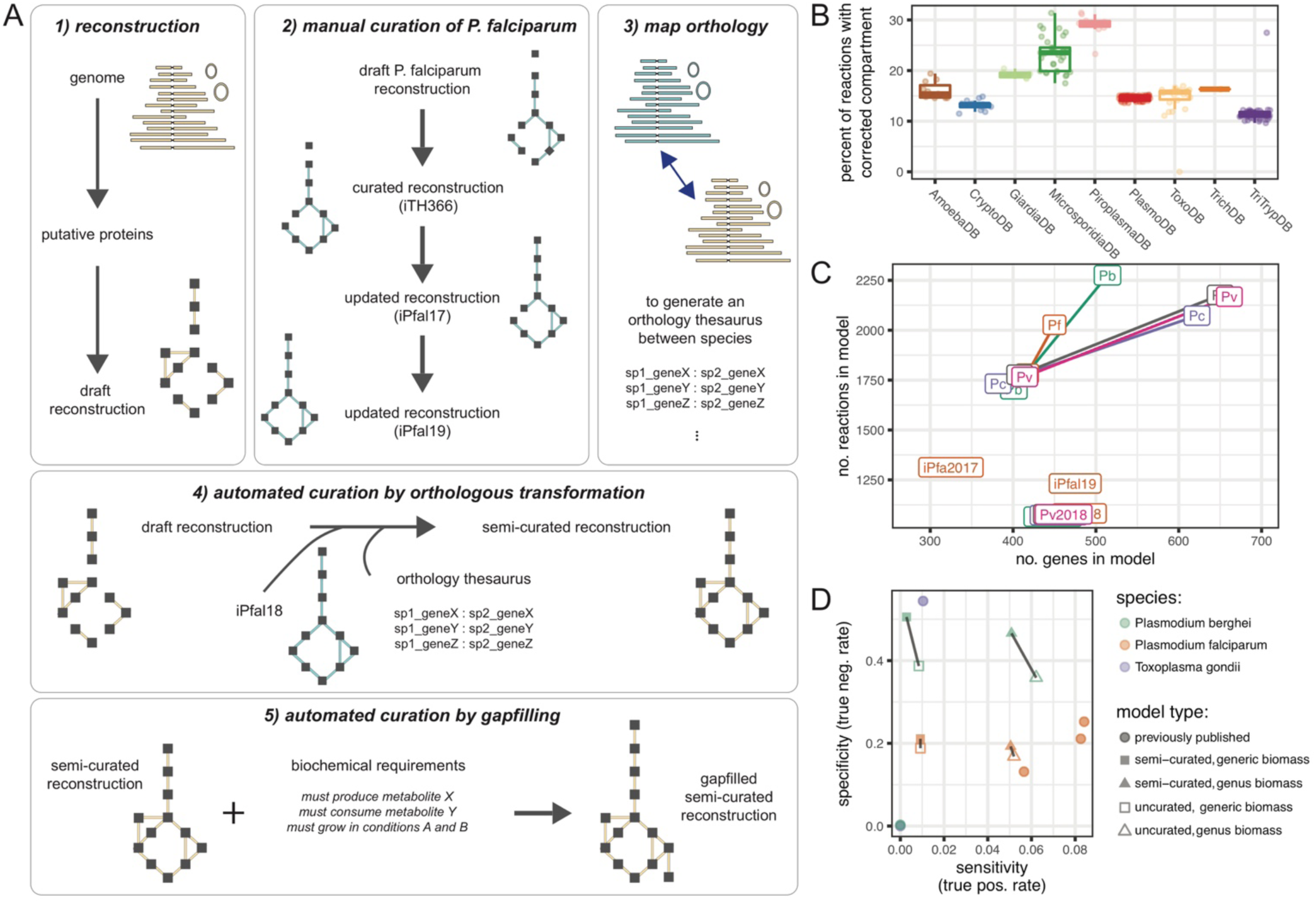
Building a parasite knowledgebase. **(A): Reconstruction pipeline.** First, *de novo* reconstructions are built from annotated genomes (see **Methods**). Next, we curated an existing manually curated reconstruction for *P. falciparum* 3D7. Third, we mapped orthologous genes so that (fourth) we could add all metabolic functions from our curated *iPfal19* into the *de novo* reconstruction by transforming each gene-protein-reaction rule via orthology. Lastly, we performed automated curation by gapfilling reconstructions to known metabolic capabilities and to generate biomass. **(B): Considering compartmentalization.** Our approach moves a large proportion of the reconstruction’s reactions from compartments in a biochemical database to biologically-relevant compartments (*e.g.* periplasm to extracellular). Thus, our *de novo* reconstruction approach accounts for compartmentalization, unlike previous metabolic network reconstruction pipelines. Each model is represented by a point. **(C): Orthology adds information.** Orthology-based curation improves reconstruction scope regarding total number of genes and reactions. These semi-curated reconstructions are larger in scope due to the addition of reactions associated with genes added via orthologous-transformation. Species are abbreviated to the genus and species initials. Pb = *P. berghei*, Pc = *P. cynomolgi*, Pf = *P. falciparum*, Pk = *P. knowlesi*, Pv = *P. vivax*. Longer names indicate previously published reconstructions (iPfal19, from Untaroiu et al. (2019) and Carey, Papin, and Guler (2017), iPfa2017 from Chiappino-Pepe et al. (2017), all others from Abdel-Haleem et al. (2018)). **(D): Prediction accuracy.** Semi-curated reconstructions recapitulate the biology of experimentally-facile parasites as well as published, manually-curated reconstructions. We tested accuracy of model predictions from the *de novo* reconstruction, the orthology-translated reconstruction, and the final semi-curated reconstruction for *P. bergehi* and compared these summary statistics to the prediction accuracy generated by our well-curated *iPfal19* and other published reconstructions (Chiappino-Pepe et al. 2017; Abdel-Haleem et al. 2018; Tymoshenko et al. 2015). This comparison was used to motivate our approach over *de novo* reconstruction building as our pipeline generates a reconstruction with greater predictive accuracy than *de novo* reconstruction and comparable to a well-curated reconstruction.

We next leveraged the manual curation in one parasite reconstruction (**Figure 2A**, *step 2*, curation from (Carey, Papin, and Guler 2017; Untaroiu et al. 2019) and in **Supplemental Table 1**) to generate a semi-curated reconstruction for a subset of phylogenetically-related organisms. To build these semi-curated reconstructions, we transformed the manually-curated reconstruction using genetic orthology (**Figure 2A**, *step 3*) and added all transformed reactions to the recipient *de novo* reconstruction (**Figure 2A**, *step 4*). Lastly, all draft and semi-curated reconstructions were gapfilled using parsimonious flux balance analysis (pFBA)-based gapfilling (Biggs and Papin 2017; Medlock and Papin 2018) to complete biochemical requirements identified in the experimental literature (**Figure 2A**, *step 5*) and to produce biomass (see the *Supplemental Methods*). As a result, when compared to manually-curated parasite reconstructions (Carey, Papin, and Guler 2017; Abdel-Haleem et al. 2018; Chiappino-Pepe et al. 2017; Tymoshenko et al. 2015), semi-curated reconstructions are larger in scope than *de novo* reconstructions and generate predictions with comparable accuracy (**Figure 2C-D**). These reconstructions are also more compliant with community standards (Lieven et al. 2018; Carey et al. 2019) than previous reconstructions for parasites (representative examples shown in **Supplemental Material: Supplemental Documents**).

Our *de novo* draft reconstructions contain only genetically supported information (prior to gapfilling) and, unsurprisingly, reconstruction size is correlated with genome size (**Figure 3A-B**). The large genome of *Chromera velia* CCMP2878 (from CryptoDB with 31,799 ORFs and 3,064 reactions) corresponds to a reconstruction with the second most unique reactions with 58. However, even small reconstructions contain unique reactions prior to gapfilling (**Figure 3C**). In fact, 32 reconstructions contain at least one unique reaction (**Figure 3C**). 343 reactions are unique to just a single model and 34% of reactions are in fewer than 10% of models (**Figure 3D**, rare reactions in red). A core set of 44 reactions are contained in all 192 reconstructions; reactions shared by all models include functions such as glycolytic enzymes. Just 6% of reactions are in at least 90% of models (blue in **Figure 3D**). The relationship between model size and genome size is weakened following gapfilling (**Figure 3E**) and the frequency of rare reactions increases (*data not shown*). ParaDIGM can be used to tease apart the difference between unique, species-specific functions and poorly annotated functions to illuminate the uncharacterized fraction of parasite genomes. To illustrate additional examples of using this resource, we identified niche-specific functions, predicted fluxomics studies to identify divergent enzymes, and identified representative model systems for drug development.

**Figure 3:**
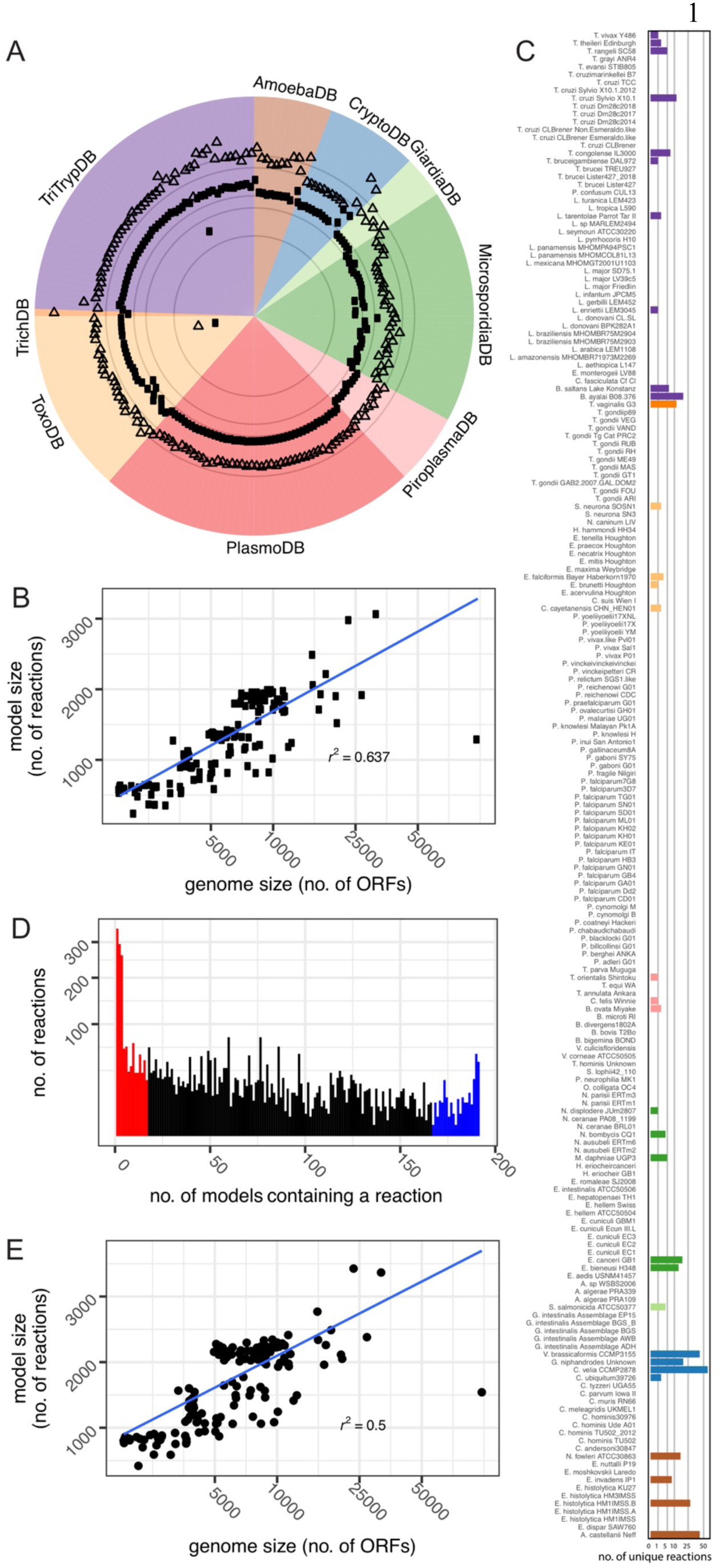
Reconstructions for all eukaryotic organisms with published genomes. **(A): Model summary.** Genome size is measured here by the number of amino acid sequences encoded by the genome (*triangle*) and model size is measured by the number of reactions present in the network (*square points*). Grey rings highlight 100, 500, 1000, 5000, and 10,000 ORFs moving from the center outwards. Genomes are grouped by database, a rough phylogenetic grouping (see **Figure 1B**). Note: *T. gondii* RH is excluded from all future analyses given its outlier nature regarding its extremely reduced genome and model size. **(B): Model size is correlated with genome size.** Larger genomes tend to generate larger models. **(C): Unique reactions.** 32 reconstructions contain at least one unique metabolic reaction, or a reaction not found in any of the other 191 models. Reconstructions are colored by EuPathDB grouping, like in panel A. **(D): Reaction frequency.** Reconstructions help identify rare metabolic functions and core parasite metabolism. **(E): Gapfilled model size remains correlated with genome size.** Following gapfilling, the relationship between genome size and model size remains; however, the correlation is weak due to an increase in the number of reactions for reconsructions built from medium-sized genomes.

### Niche-specific metabolic functions

To identify niche-specific functions, we used ParaDIGM to compare the enzymatic capacity of each organism. Specifically, we compared which enzymes are genetically supported and, therefore, present in each reconstruction prior to gapfilling. We performed classical multidimensional scaling using the Euclidiean distance between reaction presence for each reconstruction (**Figure 4A-B**). We observe that phylogenetically-related parasites tend to contain similar reactions (**Figure 4A**). However, while networks generated from genomes within a common genera or species cluster together, models also cluster within environmental niche rather than broader phylogenetic grouping such as phylum. Apicomplexan parasites cluster tightly within genus but not across genera (**Figure 4A**, Apicomplexa *in orange*). *Cryptosporidium* parasites cluster with other gut pathogens (**Figure 4A**, *in blue*) rather than other Apicomplexa. Thus, phylogeny is not the sole predictor of model similarity.

**Figure 4:**
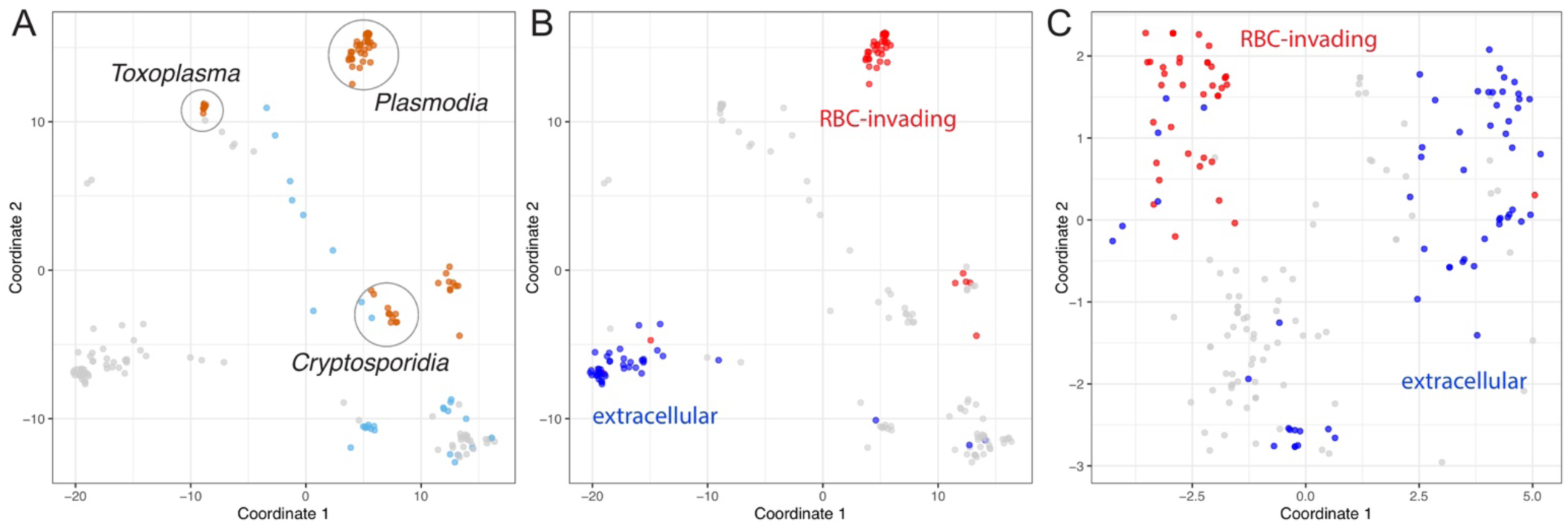
Identifying metabolic niches. **(A): Reaction content.** Classical multidimensional scaling was performed on the reaction content of all *de novo* reconstructions; thus, this analysis focuses exclusively on the genetically supported features of each reconstruction. Apicomplexan parasites (*orange*, some are circled) and all other gut pathogens (*light blue*) are highlighted. **(B): Reaction content with alternative color scheme.** Parasites that invade red blood cells (*red, Plasmodium* and *Babesia*) or can replicate extracellularly (*blue*) are highlighted. **(C): Transporter profile.** Again, parasites that invade red blood cells (*red*) or can replicate extracellularly (*blue*, like the kinetoplastids and *Giardia*, among others) are highlighted. Red blood cell-invading parasites cluster closely.

Similarly, parasites that invade red blood cells, including *Plasmodium* and *Babesia*, are dissimilar when comparing their full reaction content (**Figure 4B**, *in red*); however, the same analysis limited to each organism’s genetically encoded transporters reveals that these parasites have very similar transporter capabilities (**Figure 4C**, *in red*). This result indicates that these red blood cell-invading parasites rely on similar nutrients from their host red blood cell. On the contrary, the broad metabolic niche of extracellular growth yielded little similarity in enzymatic capacity or transporter profile (**Figure 4B** and **C**, *in blue*), likely due to the range of environments that parasites capable of extracellular growth encounter.

### Predicting metabolic function

Beyond the direct comparison of enzyme presence, we can use ParaDIGM to predict metabolic functions and the functional consequences of reaction presence and network connectivity. This approach augments the analysis beyond mere genetic comparisons: some enzymes may not be discovered in the genome despite being necessary to perform biochemical function observed experimentally and are included in these models (**Figure 5A**). Relatively few fluxomics or controlled biochemical studies have been conducted for any one organism but these data can be predicted *in silico*. We simulate fluxomics studies to profile the metabolic capability of an organism using both genetic evidence and inferred network structure. To do this, we identified which metabolites can be consumed or produced in each model following gapfilling in a rich *in silico* media, simulating the host environment. This environment is simulated by permitting import of any metabolite for which there is a genetically-encoded transporter or gapfilled transport reaction. A schematic for each metabolite categorization is shown in **Figure 5B** with experimental data shown in **Figure 5C** and analogous *in silico* results in **Figure 5D**. All models except for one (*Chromera velia* CCMP2878 with the largest genome) required gapfilling to synthesize one or more metabolites (observed experimentally) or biomass. We can expand the *in silico* predictions to all metabolites in all models (a total of 5,141 metabolites by 192 models, **Figure 5E**) to generate hypotheses about understudied metabolites and enzymes.

**Figure 5:**
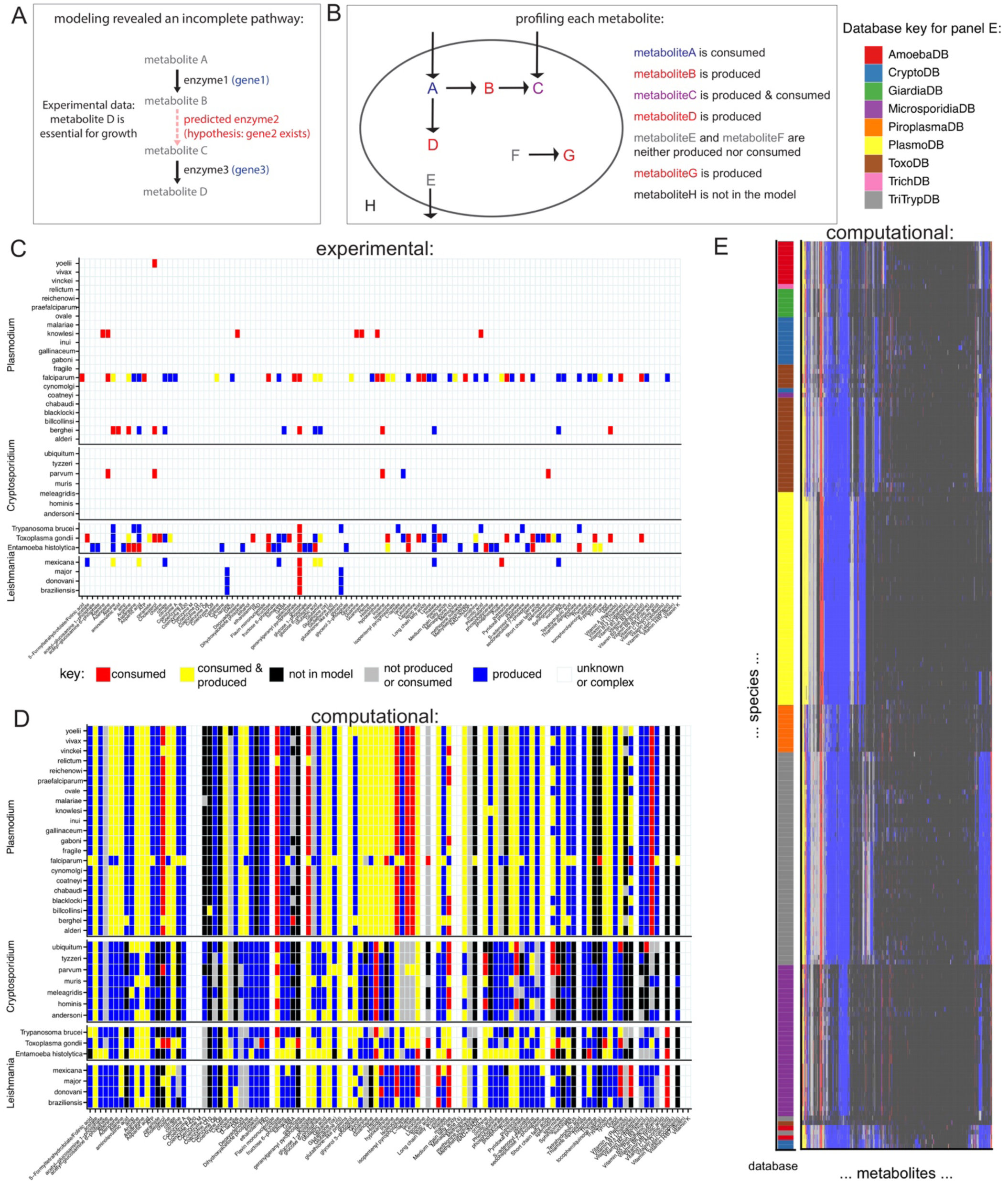
Predicting metabolic function. **(A): Advantage of network-based approaches.** Metabolic models include hypothetical functions (*i.e.* the enzyme encoded by *gene2*) that are unsupported by direct genetic evidence but may be indirectly required based on biochemical evidence. These functions are added through gapfilling. Using models augments our analysis beyond mere genetic comparisons: some enzymes may not be discovered in the genome despite being necessary for biochemical observations made and are included in these models. **(B): Defining metabolic capacities.** With our gapfilled models, we can identify if metabolites are consumed and/or produced. **(C): Experimentally-derived metabolic functions.** We compiled data providing evidence for consumption or production of select metabolites from the literature (**Supplemental Table 1**). Consumed metabolites are imported by the parasite from the extracellular environment (*e.g.* the *in vitro* growth medium). Produced metabolites are synthesized by the parasite even when the metabolite is not in the extraceullar environment. See *Online Methods* for more detail. Data are sparse. **(D): Analogous *in silico* metabolic capacity.** Inferred metabolic capacity of each organism from Panel C for every metabolite from Panel C. See Panel B for definitions. Metabolites that are neither produced nor consumed are consumed intracellularly but are not taken up from the extracellular environment. Metabolites noted as ‘complex or unknown’ here are represented by multiple metabolite identifiers in the reconstructions. **(E): Complete *in silico* metabolic capacity.** Inferred metabolic capacity of each organism (rows, clustered by database) for metabolites (columns) for every reconstruction and metabolite in ParaDIGM (5,141 metabolites by 192 models). See Panel B for definitions.

Interestingly, several metabolic enzymes were consistently predicted to be necessary for observed metabolic capabilities (**Figure 5C**) or growth across all parasites, including pyridoxal oxidase (*data not shown*). Pyridoxal oxidase is an understudied molybdenum-dependent enzyme involved in Vitamin B6 metabolism; only 24 articles on PubMed describe the enzyme, with only 3 published since 2000. Not surprisingly given the lack of literature, the reaction associated with pyridoxal oxidase is in just four reconstructions in the BiGG database, including two iterations of the *S. cervisiae* S288C model (King et al. 2016). Pyridoxal oxidase was only added to the *V. brassicaformis* CCMP3155 and *G. niphandrodes* reconstructions in the bioinformatics-driven model construction steps. However, this enzyme was added in 163 gapfilling solutions to satisfy experimentally-derived functions; thus, we predict that it is important for parasite growth. We also predict that the unidentified sequences for pyridoxal oxidase are highly divergent from known sequences because they were not identified using bioinformatic annotation methods. Thus, by comparing the reconstructions within ParaDIGM, we can identify high-confidence reactions that are encoded by divergent genetic sequences and missed by purely bioinformatic approaches.

### Selecting the most representative model system for an experiment

Genome-wide essentiality screens are available for *Plasmodium falciparum* (Zhang et al. 2018) and *berghei* (Bushell et al. 2017), *Toxoplasma gondii* (Sidik et al. 2016), and *Trypanosoma brucei*. Using the models generated with ParaDIGM, we can perform the equivalent *in silico* simulations regardless of experimental genetic tractability (**Figure 1A**, *in silico* results in **Figure 2D** and *Supplemental Table 3*). These analyses can be used to identify drugs for repurposing or the best model system for testing a novel drug target. To do this, we sequentially removed each reaction from the reconstruction to identify which reactions are necessary for growth (*i.e.* production of biomass). These simulations are performed in an unconstrained model (*i.e.* all metabolites with a transporter can be imported, all enzymes can be used) to simulate the parasite’s growth intracellularly in the nutrient-rich host cell. Dissimilarity of reaction essentiality was calculated using the Euclidean distance (**Figure 6**).

**Figure 6:**
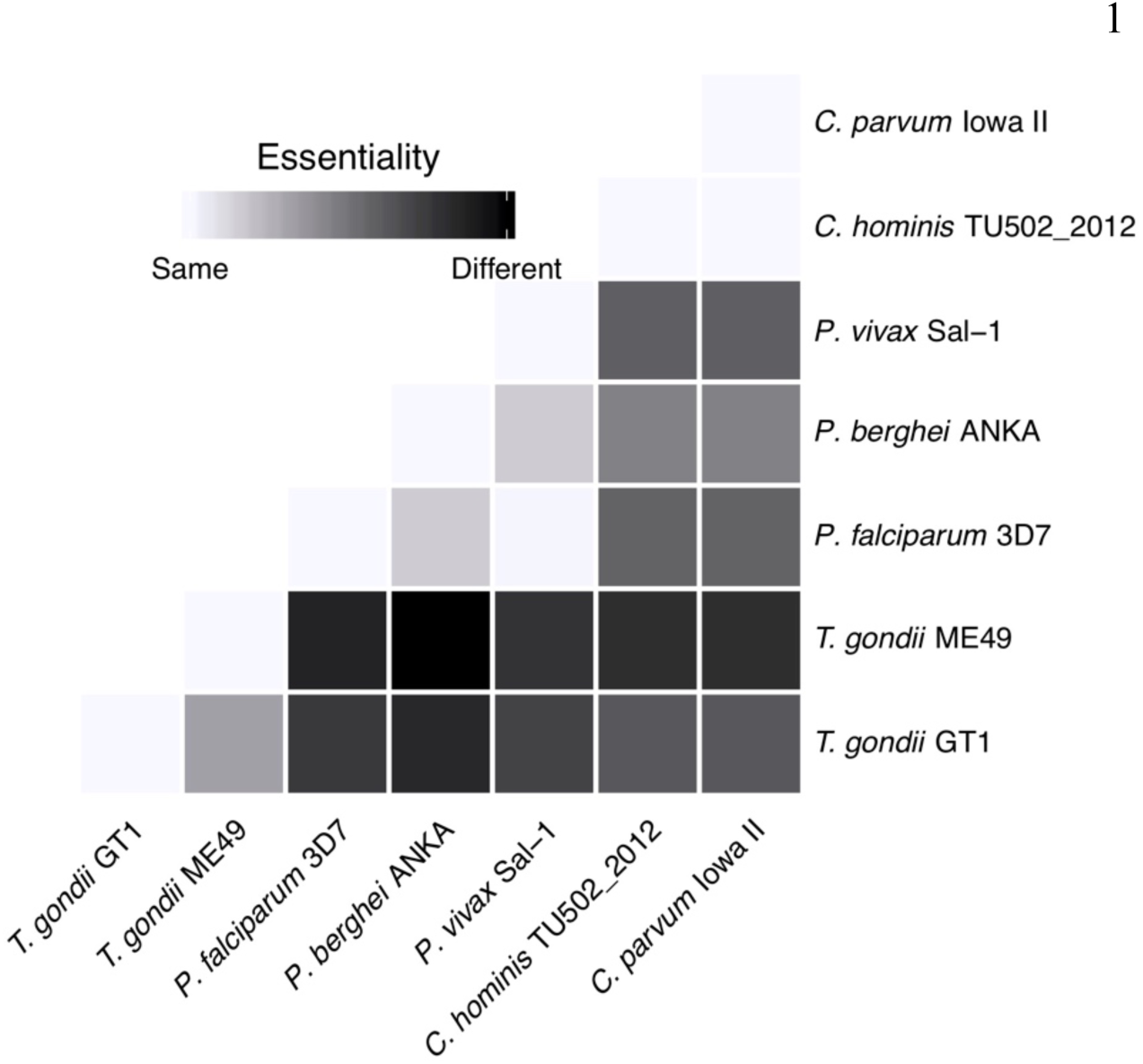
Selecting experimental model systems using reaction essentiality. Single reaction knockouts were performed on unconstrained models to identify the reactions that are essential for generating biomass. Similarity scores were calculated from binary essentiality results using Euclidean distance (root sum-of-squares of differences). *C. parvum* Iowa II and *C. hominis* TU502 (resequenced in 2012) are most similar, followed by *P. vivax* Sal-1 and *P. falciparum* 3D7. Genome-wide reaction essentiality is more similar between *Toxoplasma* and *Cryptosporidium* than *Toxoplasma* and *Plasmodium*.

While reaction essentiality is generally more similar for closely related organisms, the essentiality prediction in *T. gondii* is more similar to *Cryptosporidium* parasites than to *Plasmodium* (**Figure 6**). As *T. gondii* is a popular model system for other parasites, this result supports the use of *T. gondii* to test hypotheses about *Cryptosporidium*. Interestingly, the within-species variation in essentiality predictions for *T. gondii* is greater than across-species variation in predictions for both the *Plasmodium* and *Cryptosporidium* genera. The *T. gondii* genomes (∼63 Mbp) are much larger than the *Plasmodium* genomes (18-23 Mbp) or *Cryptosporidium* genomes (both 9 Mbp) and therefore may support greater potential for divergence. This result highlights how ParaDIGM can be used to identify functional similarities and differences between parasites that directly inform experiments for developing and studying new drugs.

## Discussion

Here, we presented a novel pipeline for generating high-quality metabolic network reconstructions from eukaryotic genomes and applied it to create 192 reconstructions for parasites. These reconstructions represent the first genome-scale metabolic network reconstructions for many of these organisms, making ParaDIGM the broadest computational biochemical resource for eukaryotes to date. ParaDIGM uses reaction and metabolite nomenclature from the Biochemical, Genetic and Genomic knowledge base (BiGG, which includes both microbial and mammalian genome-scale metabolic network reconstructions) (King et al. 2016), facilitating future work involving host-pathogen interaction modeling. Gene nomenclature used in ParaDIGM is from the Eukaryotic Pathogens Database (EuPathDB) (Aurrecoechea et al. 2017), consistent with the parasitology field standards and ‘omics data collection. Reproducible data integration approaches are used to curate each reconstruction; code and data are available in the **Supplemental Material**. ParaDIGM or individual reconstructions can be used for comparative analyses or applied to interrogate clinically- and biologically-relevant phenotypes. The adherence to community standards for metabolic modeling throughout ParaDIGM enables easier manual curation for users interested in studying a specific parasite in more detail. Together, this adherence to standards and the automated approach for integration of experimental data, will accelerate further curation of ParaDIGM itself as genome annotation improves, more experiments are performed with individual parasites, and ParaDIGM users provide feedback on reconstruction usage and performance.

This eukaryote-specific reconstruction process (**Figure 2A**) generates high-quality, comprehensive networks (**Figure 2B-D**). However, manual network curation remains the gold standard to maximize the accuracy of network predictions. Even so, our semi-automated curation approach enhances the genome-wide coverage of each reconstruction (**Figure 2C**) and generates models with comparable accuracy to manually curated reconstructions (**Figure 2D**). To evaluate these networks, we compare *in silico* predictions to experimental results; all have imperfect accuracy regarding gene essentiality (**Figure 2C**), emphasizing how challenging it is to make a truly predictive model without integrating extensive experimental data. High rates of false positives (when the model incorrectly identifies a gene as essential) are a product of the model building process; these reconstructions are built to summarize *all* metabolic capabilities of the organism, not the specific stage-dependent phenotype of an organism in the experimental system. Thus, constraining a reconstruction with *in vitro* expression data will reduce the false positive rate (*e.g.* Carey, Papin, and Guler (2017)).

We used ParaDIGM to better leverage model systems for drug development by identifying divergent or conserved metabolic pathways between select human pathogens. Network structures were quite unique with only 25.8% of all reactions in more than 50% of the reconstructions (**Figure 3D**); network topology did however cluster by genus, and transport ability is associated with specific host environments (**Figure 4**). Despite these structural similarities, minor topological differences in networks confer key metabolic strengths or weaknesses (**Figure 5 & 6**). We compare metabolic reaction (or enzyme) essentiality to identify the best *in vitro* system or non-primate infection model of disease for drug development (**Figure 6**). For example, enzyme essentiality is broadly more consistent between *Toxoplasma gondii* and *Cryptosporidium* parasites than between *T. gondii* and the malaria parasites. By leveraging network context (**Figure 5A & B**), we can impute fluxomic studies in all 192 parasites (**Figure 5D**) to contextualize the variable results across species in relatively few *in vitro* fluxomics studies (**Figure 5C**) and to expand these observations to untested organisms and metabolites (**Figure 5E**).

Beyond our use cases of ParaDIGM, the pipeline and reconstructions presented here can be used broadly by the field. The study of microbial pathogens generated paradigm-shifting results in biology. The study of viruses revealed basic cellular machinery present nearly ubiquitously in eukaryotic cells, such as the discovery of alternative RNA splicing in adenovirus (Berk 2016). The study of bacteria has provided a nearly real-time observation of evolution, allowing researchers to perform hypothesis-driven evolutionary biology experiments in addition to observational research (Baym et al. 2016). These microorganisms have shed light on cell biology and the history of life in impactiful yet highly unanticipated ways; experimental challenges associated with parasites have slowed their utility in this regard. However, both the genetic ‘dark matter’ of eukaryotic parasites and known parasite-specific functions are abundant; thus, parasites too have the capacity to inform our understanding of life. The reconstructions in ParaDIGM can be used broadly to contextualize existing experimental data and generate novel hypotheses about eukaryotic parasite biochemistry as it relates to the rest of the tree of life.

ParaDIGM provides a framework for organizing and interpreting knowledge about eukaryotic parasites. The reconstruction pipeline designed for ParaDIGM implements and builds on field-accepted standards for genome-scale metabolic modeling and the latest genome annotations in the parasitology field; moreover, it is uniquely tailored to eukaryotic cells by recognizing the importance of compartmentalization and the design of the objective function. The pipeline can be implemented with other organisms and re-implemented iteratively to incorporate novel genome sequences, biochemical datasets, genome annotations, and reconstruction curation efforts. The genome-scale metabolic network reconstructions organized in ParaDIGM also can be used broadly by the scientific community, using the reconstructions as-is as biochemical and genetic knowledgebases or as draft reconstructions for further manual curation to maximize the utility and predictive accuracy of the models. These reconstructions can be used to generate targeted experimental hypotheses for exploring parasite phenotypes, ultimately improving the accessibility of modeling approaches, increasing the utility of parasites as model systems, and accelerating clinically-motivated research in parasitology.

## Acknowledgements

The authors would like to acknowledge the helpful discussion and feedback from members of the Petri, Mann, Guler, and Papin labs, as well as Drs. Alison Criss, Norbert Leitinger, Herve Agaisse, and Young Hahn. The authors would also like to thank the University of Virginia ARCS staff for support regarding UVA’s High Performance Computing cluster, especially Karsten Siller and Katherine Holcomb. Lastly, the authors would like to thank the EuPathDB, BiGG, and CobraPy communities for providing essential tools (software and database infrastructure) and inspiration.

## Author contributions

MC conceived of the project, designed the project and analyses, and wrote the manuscript. GM and JP contributed to data analysis. GM led early adaptation of the pipeline to the University of Virginia’s High Performance Computing cluster. MS performed lipid metabolism curation. GM, WP, JG, and JP provided feedback and suggestions which helped shape the project. All authors also edited and approved the manuscript.

## Supplemental Information

### Code, data, and reconstruction availability

All code, data, and resultant reconstructions are available at https://github.com/maureencarey/paradigm.

## Online Methods

### Code dependencies

R (R Core Team 2017) and R packages tidyverse, ggpubr, ggdendro, seqinr, Biostrings, msa, reshape2, UpSetR, cluster, ade4, RColorBrewer, readxl, dplyr, and ggdendro were used for analysis or visualization (Hadley Wickham 2017; Kassambara 2017; Vries and Ripley 2013, 2013; Charif and Lobry 2007; Pagès et al. 2017; Bodenhofer et al. 2015; Wickham 2012; Gehlenborg 2017; Maechler et al. 2013; Dray, Dufour, and Others 2007; Neuwirth and Brewer 2014; Hadley Wickham and Bryan 2017; Wickham et al. 2015). Python 3.6.4, pandas, and CobraPy 0.14.1 (Ebrahim et al. 2013), as well as code to implement Diamond-based annotation scoring from CarveMe (Machado et al. 2018), were used for the reconstruction and modeling. Memote (Lieven et al. 2018) was used to evaluate all reconstructions.

### Genomic Analyses

Sequences were obtained from EuPathDB release 44 (Aurrecoechea et al. 2017). EuPathDB curates and compiles genome annotation for all genomes hosted by the database. We used open reading frames identified on EuPathDB and annotated the sequences with Diamond, described below. EuPathDB’s OrthoMCL was used for mapping orthology between *Plasmodium* species. In brief, orthology was mapped within each EuPathDB database by the ‘map by orthology’ tool from the genome of each organism with a curated reconstruction to all other genomes within that database. The search protocol was ‘new search > genes > taxonomy > organism [pick] > transform by orthology’. We mapped each organism’s amino acid sequences using Diamond annotation (Buchfink, Xie, and Huson 2015) against proteins referenced in the BiGG databases (King et al. 2016) or against protein sequences obtained from OrthoMCL, part of EuPathDB that contains orthologous groups of parasite genes (Li, Stoeckert, and Roos 2003). Diamond is a similar approach to BLAST, with sensitive and fast performance on protein annotations (Buchfink, Xie, and Huson 2015).

### Model Generation

We generated draft reconstructions by first annotating each organism’s amino acid sequences, obtained from EuPathDB (Aurrecoechea et al. 2017), using Diamond annotation (Buchfink, Xie, and Huson 2015) against proteins referenced in the BiGG databases (King et al. 2016). We next mapped all functional annotations to reactions contained in the BiGG database (King et al. 2016) inspired by the approach conducted with the reconstruction pipeline CarveMe (Machado et al. 2018). Methods are included in the analytic code hosted on Github, at https://github.com/maureencarey/paradigm.

Unlike the CarveMe approach (Machado et al. 2018), we included all high-scoring reactions. CarveMe maximizes the number of high-scoring hits while building a functional network. Our conservative approach generates broadly inclusive but incomplete reconstructions (*i.e.* that are not able to produce biomass until gapfilled). This approach added redundant reactions from multiple different compartments (*e.g.* peroxisome, mitochondria, and cytosol) so all reaction versions other than the cytosolic version were pruned unless localized to a relevant compartment (**Supplemental Table 2**); for genera not included in **Supplemental Table 2**, only the cytosol and extracellular space were used. For example, if a *Plasmodium* reconstruction contained a reaction in the cytosol, mitochondria, and chloroplast, only the cytoplasmic and mitochondrial reaction versions would be kept. Following this step, a large percentage of each reconstruction’s reactions remained in unsupported compartments because there was no analogous cytosolic reaction (**Figure 2**). Next, reactions only found in an unsupported compartment were moved to the extracellular space or cytosol; specifically, periplasmic metabolites were moved to the extracellular space and all internal subcompartment metabolites were moved to the cytosol. However, this step removed all reactions that summarized a transport reaction from the extracellular space to periplasm or from the cytosol to an unsupported organelle. The extracellular compartment corresponds to the parasitophorous vacuole space contained within the host cell for intracellular parasites (*i.e. Plasmodium, Toxoplasma, Cryptosporidium*) and the host serum for extracellular parasites (*i.e. Trypanosoma*).

### Manual Curation

We performed brief manual curation from literature sources, building on our curation conducted in Carey, Papin, and Guler (2017) and Untaroiu et al. (2019). Our previous asexual blood-stage *Plasmodium falciparum* 3D7 reconstruction (Untaroiu et al. (2019) adapted from Carey, Papin, and Guler (2017)) was manually curated generating *iPfal19*. See supplemental information for code documenting all for modifications and implementation of manual curation.

Additional manual curation was performed on lipid metabolism of the asexual blood-stage *P. falciparum* using the lipidomics study presented in Gulati et al. (2015) adding over 700 reactions, 400 metabolites, and 18 genes. This curation removed aggregate reactions representing lipid metabolism and replaced them with individual reactions for individual lipid species, as supported by the lipidomics study. This model is available, but was not used for the analyses presented here as the metabolic demands for lipids in our biomass reaction are also aggregated. Inclusion of these reaction is appropriate for understanding lipid metabolism but would create random distributions of flux through the individual reactions that may distract from meaningful changes in flux distributions. All code for this curation is available at https://github.com/gulermalaria/iPfal17_curation.

### Automated orthology-driven curation

We developed a novel automated curation approach using orthologous transformation, similar to the approach taken by Abdel-Haleem et al. (2018). Our approach leverages the curation conducted in one organism for closely-related organisms and applied it to all draft *Plasmodium* reconstructions using *iPfal19* (**Figure 2A**). We first mapped orthology of *P. falicparum* to each other *Plasmodium* species to build an orthology thesaurus (**Figure 2A**). We then added genes and associated reactions from *iPfal19* if there was an orthologous gene in the target species’ reconstruction (**Figure 2A**) resulting in a significant increase in the number of genes and reactions in each reconstruction (**Figure 2C**). Notably, this approach facilitates the compartmentalization of these reconstructions, a function many automated pipelines fail to include. This step is particularly important for parasite-specific compartments such as the apicoplast, which is not included in any database.

### Gapfilling (part 1) - Automated data-driven curation

Gapfilling is an analytic process used to bridge or complete genetically-supported metabolic pathways to permit the network to fulfill metabolic functions, and was used to generate functional models. To increase the scope of a reconstruction (*i.e.* to add reactions), we performed gapfilling to fill in gaps in pathways to ensure that the reconstruction can complete a particular task. This optimization process adds reactions to allow the reconstruction to carry flux under given constraints.

Further automated curation of all reconstructions was performed by gapfilling for metabolites measured to be consumed in fluxomic or select media formulation studies. Detailed analysis is provided in our code, Following an extensive literature review, we compiled data providing evidence for consumption or production of select metabolites (**Supplemental Table 1**). Metabolites were defined as consumed by the parasite if: (1) the metabolite was radiolabeled, added to media, incorporated into the parasite or converted by the parasite, and this was not seen to the same degree in uninfected host cells; (2) the metabolite rescued inhibitor treatment of a metabolically upstream parasite enzyme; or (3) the metabolite is an essential media component for parasite culture. Metabolites were defined to be produced by the parasite if the metabolite was radiolabeled following growth in a media containing a radiolabeled precursor metabolite, and this was not seen to the same degree in uninfected host cells. First, import or excretion of these metabolites were added to the reconstruction. Next, the model objective was changed to an internal demand reaction for the metabolite or excretion reactions, respectively, and was gapfilled sequentially; this ensures import or synthesis of each of these measured metabolites.

To gapfill for individual metabolites or biomass (next section), we used a parsimonious flux balance analysis (pFBA)-based approach as originally used in Biggs and Papin (2017) and futher developed in Medlock and Papin (2018). Code is linked in the Supplemental Information. In essence, this pFBA approach minimizes the flux through genetically unsupported reactions from a biochemical database such that the network can carry flux through the objective reaction (*i.e.* metabolite production or consumption or biomass synthesis). Any reaction from the database that carries flux during this problem was added to the network.

### Gapfilling (part 2) - Biomass as the objective function

After compartmentalization, manual curation, and gapfilling for individual metabolites, we used pFBA-based gapfilling to ensure each network was capable of generating biomass. We use two classes of biomass functions here to robustly evaluate model performance, specifically a genus-specific curated biomass reaction (for *Plasmodium* reconstructions) and a parasite-specific generic biomass reaction. The genus-level curated biomass reaction was taken from our manually curated *Plasmodium falciparum* model. Our generic biomass contains metabolites from several curated reconstructions and thus contains metabolites from the *Plasmodium falicparum, Leishmania major*, and *Cryptosporidium hominis* species-specific biomasses with the stoichometry contained in the *iPfal19* biomass reaction. Unfortunately, variability in reconstruction namespace (*i.e.* the database used for metabolite and reaction nameing conventions) make it difficult to access data compiled for some parasite reconstructions, such as the *Toxoplasma* and *Plasmodium* reconstructions *ToxoNet1* and *iPfal19*, respectively, as there are not always one-to-one mappings of variables across databases. This generic biomass was used to capture the most conservatively defined required *parasite* biosynthetic capacity. All reconstructions were gapfilled to ensure biomass could be synthesized via the generic parasite biomass reaction; all *Plasmodium* reconstructions were also gapfilled to ensure biomass could be synthesized via the genus-specific biomass reaction.

### Model Performance Evaluation

Network accuracy was evaluated against gene essentiality data (**Supplemental Table 3**) if available. Gene essentiality was simulated by performing single gene deletion studies in our models. All simulations were performed in a model state without additional experimental constraints; specifically, all exchange reactions were permitted to carry flux reversibly, simulating a nutrient rich environment such as intracellular growth. Of note, these ‘unconstrained’ models do not necessarily represent the *in vitro* or *in vivo* environment in which all experiments were conducted, merely the metabolic capacity an organism encodes. If this extracellular environment was constrained to represent the specific host environment, we would see further separation of model predictions.

Gene deletions were simulated by removing the gene of interest from the model using CobraPy’s ‘single_gene_deletion’ function. This change results in the inhibition of flux through all reactions that require that gene to function. If the model could not produce biomass with these constraints, the gene was deemed essential. Specifically, we defined an essential gene as a knockout that resulted in 10% or less of the maximal biomass (measured in mmol/(gDW*hr)). Knockout accuracy was defined as the sum of true positives (refractory to knockout or lethal genes) and true negatives (nonlethal genes) divided by the total number of predictions. As targeted metabolomics data were used for model generation (by gapfilling), we excluded these data from the evaluation data.

Models were tested for thermodynamically-infeasible loops and energy-generating cycles; the approach outlined in Fritzemeier et al. (2017) was used with minor modifications for eukaryotic cells.

### Memote Evaluation

Models were evaluated using Memote (Lieven et al. 2018); example outputs are presented in the **Supplemental Material: Supplemental Documents**. Additionally, Memote was used to quality control the reconstructions throughout the development of the ParaDIGM pipeline to improve standard compliance (especially annotation coverage) and biological relevance (*e.g.* network connectivity and topology, *data not shown*).

### Supplemental discussion of our reconstruction pipeline for eukaryotes

Our pipeline itself (**Figure 2A**) is uniquely tailored to eukaryotic cells by recognizing the importance of compartmentalization and the design of the objective function. Compartmentalization biases predictions (Carey, Papin, and Guler 2017); both our *de novo* reconstruction and orthology-driven approaches addresses this important step. Compartmentalization was incorporated into our *de novo* reconstruction pipeline and implemented for several genera (**Supplemental Table 2**). We adjusted the localization of reactions if inappropriately compartmentalized and added unique parasite-specific compartments (*e.g.* the apicoplast in *Plasmodium*). This addition was done by adding genetically supported reactions were to all feasible compartments. If a gene-encoded enzyme corresponds to both mitochondrial and cytoplasmic reactions, both versions will be included: adding network redundancy that may not be biologically accurate. Alternatively, if an enzyme maps to a chloroplast reaction, our approach moved the reaction to the cytosol. It is plausible that chloroplast reactions like this example are not catalyzed by the parasite. These modifications are encoded in our analytic pipeline for future reference (see code, **Supplemental Information**). Additionally, we used a curated model to inform the compartmentalization of each semi-curated model (**Figure 2A**); genes associated with compartmentalized reactions were mapped via orthology, assuming orthologous genes has comparable localization across species.

Similarly, the objective function (often representing biomass synthesis (Feist and Palsson 2010)) influences network simulations (Xavier, Patil, and Rocha 2017) and the assumptions used in formulating a biomass reaction for prokaryotes may not apply to eukaryotes (Liebermeister et al. 2014). For example, in the first genome-scale metabolic model of any *Cryptosporidium* species, *C. hominis* (Vanee et al. 2010), 30 reactions involved in lipid synthesis were unsupported by genetic evidence and manually added to the network. The selection of lipid species as biomass precursors may have impacted the addition of these unsuported reactions and resultant simulations with the model; for example, the 30 gapfilled reactions might not be added if alternative or fewer lipid species were included in the biomass reaction.

Thus, we use a consensus biomass reaction derived from multiple curated parasite reconstructions, as well as a genus-specific biomass for *Plasmodium* reconstructions. These objective functions influence reactions added via gapfilling, adding uncertainty in network structure (Biggs and Papin 2017; Medlock and Papin 2018); thus, we emphasize which reactions were gapfilled frequently (*i.e.* for many species within a genera or ParaDIGM broadly) to increase our confidence in these predicted functions. Targeted investigation of these reactions (such as pyridoxal oxidase and shikimate dehydrogenase) will increase our confidence in all of our parasite network reconstructions.

## Supplemental Materials

**Supplemental Table 1:Automated curation tasks.** All reconstructions were gapfilled to ensure the network could consume or produce all relevant metabolites outlined here. Data from multiple strains of one species were aggregated. This file is too large to include in this document but is available as a supplemental document and is available as: https://github.com/maureencarey/paradigm/data/auxotrophies_references.xlsx.

**Supplemental Table 2:** Compartmentalization. Subcellular compartments found in each genus.

| Species/Database | Compartments |
| --- | --- |
| BiGG database | cytosol, extracellular, mitochondria, nucleus, lysosome, chloroplast, golgi, vacuole, endoplasmic reticulum, peroxisome/glyoxysome, flagellum, periplasm, thylakoid, thylakoid membrane, cytosolic membrane, carboxyzome, intermembrane space of mitochondria, eyespot, unidentified |
| Plasmodium | cytosol, extracellular, mitochondria, apicoplast, food vacuole |
| Leishmania | cytosol, extracellular, mitochondria, kinetoplast, glycosome |
| Cryptosporidium | cytosol, extracellular, pseudomitochondria |
| Toxoplasma | cytosol, extracellular, mitochondria, apicoplast |
| Giardia | cytosol, extracellular |
| Entamoeba | cytosol, extracellular |
| All else | cytosol, extracellular |

**Supplemental Table 3:** Available essentiality datasets. Data available for evaluating parasite gene essentiality.

| Organism | Dataset type | Reference (DOI) |
| --- | --- | --- |
| *Toxoplasma gondii* GT1 | CRISPR site-directed mutagenesis | Sidik 2016 (10.1016/j.cell.2016.08.019) |
| *Plasmodium falciparum* 3D7 | piggyBac transposon mutagenesis | Zhang 2018 (10.1126/science.aap7847) |
| *Plasmodium berghei* ANKA | transfection mutagenesis | Bushell 2017 (10.1016/j.cell.2017.06.030) |
| *Trypanosoma brucei* 2T1 | RNAi screen | Stortz 2017 (10.1371/journal.ppat.1006477) |
| *Trypanosoma brucei* MiTat 1.2, clone 221a | RNAi screen | Alsford 2011 (10.1101/gr.115089.110) |

**Supplemental Documents: Memote reports for iPfal19 and three ParaDIGM reconstructions,** *P. falciparum Dd2, T. gondii* ME49, and *C. tyzzeri*, are available at https://github.com/maureencarey/paradigm/memote_reports.

